# Investigating the mechanisms of indocyanine green (ICG) cellular uptake in sarcoma

**DOI:** 10.1101/2021.04.05.438013

**Authors:** Corey D Chan, Marcus J Brookes, Riya Tanwani, Chloe Hope, Toni A Pringle, James C Knight, Kenneth S Rankin

## Abstract

**Introduction:** Indocyanine green (ICG) is a near infrared (NIR) dye which has been used clinically for over 50 years and has recently been utilised for fluorescence guided surgery in a number of cancer types, including sarcoma. ICG is taken up and retained by sarcoma tumours to a greater extent than normal tissue, demonstrating its potential to aid in visualisation of tumour margins. However, the mechanisms surrounding preferential ICG uptake in tumours are poorly understood.

**Methods:** *In vitro* ICG cellular uptake studies were performed across a panel of four sarcoma cell lines and one breast cancer cell line, exhibiting varying proliferation rates and phenotypes. The effects of ICG concentration, incubation time, inhibition of clathrin mediated endocytosis and cell line proliferation rate on the cellular uptake of ICG were investigated using fluorescence microscopy and flow cytometry. The spatial orientation of ICG was also assessed in a patient specimen.

**Results:** The level of ICG cellular uptake was dependent on ICG concentration and incubation time. Cell line proliferation rate correlated significantly to ICG uptake within 30 minutes (Rs= 1, p<0.001), whilst retention of ICG after 24hrs did not (Rs= 0.3, p=0.624). From our data, the primary mechanism of ICG uptake in sarcoma cells is via clathrin mediated endocytosis. Following the resection of a grade 3 leiomyosarcoma, ICG signal was detectable macroscopically and on 3μm sections, whilst being negative on the muscle control.

**Conclusions:** The use of ICG for tumour detection in sarcoma surgery may demonstrate higher utility in high grade tumours compared to low grade tumours, due to the observation of higher ICG uptake in more proliferative cell lines. It is likely that the enhanced permeability and retention (EPR) effect plays a significant role in the retention of ICG within tumours. Future work on the detection of ICG at the cellular level within human tissue sections is required, with the aid of purpose built NIR microscopes.

## Introduction

Indocyanine green (ICG) is a near infrared (NIR) fluorescent tricarbocyanine dye^1^ with numerous uses in the medical field, mainly in angiography and fluorescence guided oncological tumour resection^2^. ICG has a well-established safety profile, with minimal side effects, and is approved for clinical use by the Food and Drug Administration and the European Medicines Agency. The main safety consideration for use in patients is a known allergy to iodine, which has been speculated to increase the risk of a systemic reaction^3^; anaphylaxis to ICG is extremely rare and has previously been reported as less that 1 in 10,000^4^.

ICG is water-soluble and has well-established absorption and emission properties^5^. At low concentrations, ICG has an absorption peak at 780nm and emission peak at 830nm; as the concentration increases, ICG has a tendency to form oligomers from its previously monomeric structure, causing the absorption peak to shift to 695nm^5, 6^. The emission properties vary even further when the solvent includes plasma or human albumin^5, 7, 8^. This is likely owing to its amphiphilic structure, with both hydrophilic and hydrophobic moieties^1^, resulting in ICG binding to lipoproteins and albumin in the blood via ICG’s hydrophilic component^9, 10^, with binding to the vascular endothelium secondary to affinity to the phospholipid bilayer also demonstrated^8^.

ICG is rapidly excreted by the biliary system, with a plasma half-life of 2-4 minutes^10^, and, as such, found early use in the measurement of hepatic function^11-13^. Over the past 50 years, ICG has also been adopted for other medical uses, including ophthalmic imaging and the monitoring of cardiac output due its predictable and extensively studied pharmacokinetic properties^14^. Its primary use has since switched to that of lymph node mapping, angiography and fluorescence-guided surgery. Intravenous administration allows quantitative assessment of tissue perfusion intraoperatively, allowing assessment of skin flap viability and congestion, prompting immediate revision, as opposed to future operations after flap failure^15-17^; ICG angiography has also been demonstrated as a bedside tool to improve detection of reduced perfusion postoperatively^18^. Besides angiography for flap perfusion, ICG has also been described during intra-cranial aneurysm surgery^19^, coronary artery bypass graft surgery^20^, and is currently under investigation to evaluate whether it can reduce the anastomotic leak rate in colorectal cancer resection^21^.

The use of ICG in cancer surgery is a rapidly progressing field, with its use for the identification of satellite lymph nodes in breast^22, 23^, gastro-intestinal^24-26^ and gynaecological malignancies^27^ well described. ICG is now being used to identify malignant tumours, rather than just the lymph nodes they drain to. ICG has been shown to accumulate in the tumours of multiple malignancy types, resulting in the area fluorescing when viewed under a near infrared camera. This was first described in liver tumours, in which both hepatocellular carcinomas and metastatic lesions have been shown to fluoresce^28, 29^. The use of ICG for fluorescence guided surgery has since been described in both breast cancer^30^, in which the absence of fluorescence in the surgical bed was predictive of negative margins^31^, ovarian cancer^32^, head and neck cancer^33^, lung cancer^34^ and sarcomas^35^. Unlike angiography, in which the ICG is given intraoperatively, the ICG is instead given pre-operatively. The optimal timing of administration is unclear, with the reported protocols varying widely; Morita et al. administered the drug up 28 days pre-operatively for the detection of hepatocellular carcinomas^36^, whilst Bourgeois et al found that administering the dye immediately pre-operatively improved tumour detection in breast cancer^30^.

Despite the use of ICG for fluorescence guided tumour resection now being well described, the mechanism by which the dye accumulates within and is retained by the tumour, is not well understood. The accumulation of ICG within the tumour has been postulated to be secondary to the enhanced permeability and retention (EPR) effect^37^; the EPR effect describes a theory in which macromolecules selectively accumulate in the tumour secondary to the leaky endothelium of the neovasculature and are retained secondary to impaired lymphatic drainage^38^. Maeda et al. theorised that, as ICG readily binds to albumin and globulin *in vivo*, it selectively extravasates only through the leaky tumour vasculature, otherwise being rapidly cleared from the plasma by the liver^37^. This was backed up by Madajewski et al who demonstrated in a mouse model that fluorescence correlated with tumour vasculature density^39^.

Work by Onda et al. contradicted this theory, demonstrating ICG to rapidly extravasate in all tissues in their mouse model of colorectal cancer, with no tumour specific delivery demonstrated^40^. They did, however, demonstrate that ICG was taken up preferentially by tumour cells via clathrin mediated endocytosis (CME) due to the high endocytic rate of cancer cells^40^, suggesting this may be related to ICG’s affinity to phospholipids and its inherent ability to bind to cell mambranes^9^. They also demonstrated that tumour cells retained the dye for a minimum of 24 hours, as opposed to normal tissues from which it was rapidly cleared. This suggested overall that the primary mechanism of tumour fluorescence was increased cellular uptake and retention as opposed to the EPR effect. However, these findings contrast somewhat with recent work by Sardar et al., who found ICG in higher relative concentration in the acellular, necrotic areas of their mouse model of synovial sarcoma^41^; the accumulation of ICG in necrotic areas of non-cancerous tissues has also previously been demonstrated^42, 43^.

With these contrasting findings, it is apparent that the mechanism of the intra-tumour accumulation of ICG is still not clear. It is highly likely that aspects of both models discussed above contribute to enhanced tumour uptake of ICG, including the intrinsic dysregulation of cancer cell pathways, as well as the tumour microenvironment and EPR effect. These factors will of course vary depending on the type of tumour and if adjuvant therapy has been utilised. An improved understanding of ICG mechanisms will allow a more informed clinical use of the dye, helping to optimise aspects of its use such as the dose, route and timing of administration, which remains unclear. In this study, our aim was to further understand the mechanism of action of ICG in sarcoma, by assessing uptake and retention across a range of sarcoma cell lines, as well as processing and imaging human fluorescing tumour tissue sections post-operatively.

## Material and Methods

### Cell Culture

Five human cancer cell lines were used in this study, including four human sarcoma cell lines (HT1080, U2OS, MG63, SAOS2) and one breast cancer cell line as a comparative control from an alternative cancer type with low proliferation (MCF-7). Cell lines were chosen to represent sarcoma cells of varying proliferation rates. HT1080 is a rapidly proliferating cell line derived from a dedifferentiated chondrosarcoma. U2OS, MG63 and SAOS2 are osteosarcoma cell lines with varying proliferation rates (SAOS2 has a slow proliferation rate). All cell lines were cultured in RPMI 1640 medium supplemented with 10% foetal bovine serum, 100 U penicillin/mL and 0.1 mg streptomycin/mL (henceforth being referred to as normal media) and incubated at 37°C in a 5% CO_2_ humidified atmosphere. For Pitstop 2 experiments, buffered serum free media was used consisting of DMEM/F12 (1:1) with 15mM HEPES, supplemented with 100 U penicillin/mL and 0.1 mg streptomycin/mL with no added foetal bovine serum. All of these cell lines are adherent but differ in their morphology and proliferation rate. Further information for these cell lines can be found in the Supporting Information (Table S1).

### Fluorescent Agents

Clinical grade ICG was obtained from Verdye in the form of vials containing 25mg lyophylised powder. ICG powder was weighed and dissolved in cell culture medium (type of medium dependent on experiment) and sterile filtered (Sartorius Minisart High Flow, PES 0.22μm) to achieve a 100μM stock solution. The stock solution was stored in the dark at 4°C and used for a maximum of 5 days. For microscopy, cell nuclei were stained with Abcam Mounting Medium with DAPI Aqueous Fluoroshield (ab104139).

### Cell preparation in chamber slides

Cells were seeded at a density between 2 × 10^4^ and 4 × 10^4^ cells per well in Nunc Lab-Tek II CC2 chamber slides (Thermo Scientific) based on their proliferation rate and left to establish for 48hrs at 37°C. Media was aspirated, and cells incubated in ICG in normal media for either 15 minutes or 30 minutes. ICG was removed, cells washed x1 with normal media then x2 with PBS, fixed with 4% formaldehyde solution (pH 6.9, buffered, Sigma-Aldrich) for 10 minutes at room temperature. PFA was removed, cells washed x3 in PBS, the chamber system was removed, the cells stained with DAPI mounting medium and cover slip added. All slides were imaged immediately after mounting.

### Fluorescence Microscopy

Slides were imaged using the Zeiss Axio Imager 1 at 20x and 40x magnification. ICG fluorescence was detected using the Cy7 cube; bandpass filter excitation 670-745, emission 768-850, exposure time 2000ms – 15,000ms. DAPI was detected using Zeiss 49 cube; bandpass filter excitation 335-385, emission 420-470, exposure time 80ms. Further details on the microscope set-up can be found in the Supporting Information.

### Image quantification

Microscopy images were quantified at 40x magnification, by calculating the mean fluorescence intensity per cell. The .CZI files were opened in ZEN 3.3 Blue Edition software (Zeiss), with all three channels overlayed (DAPI, Cy7, DIC). The polygonal tool was used to draw around each cell guided by the DIC overlay, and the arithmetic mean of Cy7 signal was generated by the ZEN software. Cells from each quadrant of the FOV were represented. An average was calculated across all measured cells. Further information can be found in the Supporting Information (Figure S4).

### Flow cytometric analysis

All five cell lines were seeded into wells of a 6 well plate at a density between 2.5 × 10^5^ and 5 × 10^5^ cells per well based on their proliferation rate and left to establish for 48hrs at 37°C. Cells were checked under the microscope to ensure consistent confluency across all cell lines (80-90%), washed with PBS, then incubated with ICG (25μM) in normal media for 30 minutes at 37°C. The ICG was removed, cells washed x3 with PBS, trypsinised and neutralised in media. The cells were centrifuged, washed in 10 mL ice cold flow buffer, centrifuged, then resuspended in 500μL flow buffer (for analysis on BD FACSCanto II; laser 635 nm (bandpass filter 780/60). The experiment was repeated on three separate occasions. Flow Buffer Formulation: 500mL PBS, 2.5mL of 0.2 mmol/L EDTA prepared from anhydrous stock (Sigma), 2.5mL of BSA 10% Stock Solution (MACS).

For the 24hr retention experiment, ICG was removed after 30 minutes, cells were washed x3 with PBS, and incubated for an additional 24 hours at 37°C in fresh normal media, then processed as above the following day. The samples were analysed using the same BD FACSCanto II acquisition setup as the previous day, and the data sets were directly compared. The experiment was repeated on two separate occasions. Additional details can be found in the Supporting Information (Figures S1-S2).

### Cellular Proliferation Rate

Cellular proliferation rate was assessed using Cell Counting Kit-8 (CCK-8, Dojindo Labatories, CK04-11) as per the manufacturer’s manual. Specifically, all cell lines were seeded at a density of 1.5 × 10^4^ cells per well in a 96 well plate in triplicate, alongside a media only control, and incubated for 48 hours. 10μL of CCK-8 solution was added to each well, the cells incubated for 2 hours and the absorbance measured at 450 nm using a microplate reader (Omega FLUOstar). Value calculated as (average CCK-8 absorbance value) – (average background value). The experiment was repeated on three separate occasions and an overall average value taken forward for analysis for each cell line.

### Clathrin Inhibition via Pitstop 2

For microscopy, HT1080 cells were prepared in chamber slides at a density of 2 × 10^4^ cells per well and left to establish for 48hrs. Old media was removed, and cells incubated in serum free media for 10 minutes prior to the experiment. Serum free media was used as per the manufacturer’s instructions (abcam) as Pitstop 2 (PS2) is sequestered by serum albumins. Cells were incubated with varying concentrations of Pitstop 2, Novel cell-permeable clathrin inhibitor (ab120687) or Pitstop 2 negative control (ab120688) diluted in serum free media for 15 minutes at 37°C; the final PS2 working concentrations had 0.1% DMSO concentration. Non-treated control wells were incubated in serum free media with 0.1% DMSO for 15 minutes at 37°C. After 15 minutes, 50μM ICG in serum free media was added into the pre-existing media to achieve a final ICG concentration of 25μM per well, and incubated for 30 minutes at 37°C. The media was removed, the cells washed x3 in PBS, and prepared as previously described. For flow cytometry, HT1080 cells were seeded in a 6 well plate at a density of 2.5 × 10^5^ cells per well and left to establish for 48hrs. Cells were treated with PS2 as described above, incubated with 10μM or 25μM ICG for 30 minutes and processed for flow cytometric analysis as previously described.

### Ethics approval and consent to participate

Appropriate informed consent for the use of human tumour specimen slides and paraffin embedded blocks was obtained and approved by the Newcastle and North Tyneside 1 Research Ethics Committee (REC Reference Number: 17/NE/0361).

### Patient tumour sections

75mg ICG was administered intravenously at induction of general anaesthesia. Following resection, the specimen was assessed with the Stryker SPY-PHI infrared camera. A tumour sample was taken from an area with intense fluorescence along with a muscle sample (Figure 6). Following image acquisition in the operating theatre, the samples were processed in a standard manner into paraffin blocks. 3μm sections were cut, placed on glass slides and stained with DAPI before hard mounting with coverslips.

### Biomolecular Imaging of tumour sections

Specimen slides were imaged using the Amersham Typhoon NIR Biomolecular Imager. Slides were scanned at a resolution of 10 microns using the Cy7 laser for ICG, and the Cy2 laser for DAPI. Image files were processed and analysed using Fiji Image J. There were 3 tissue samples per specimen slide. The polygon tool in Image J was used to draw around each section of tissue and the signal in Cy7 and Cy2 channels was quantified. Cy7 signal (ICG) of tumour vs muscle was calculated with respect to Cy2 (DAPI).

### Statistical Analysis

The correlations between variables were calculated using the non-parametric Spearman’s Rho function (2-tailed) on SPSS (V.27).

## Results

### Levels of ICG uptake in sarcoma cells is dependent on concentration and incubation time

On fluorescence microscopy, the detection of ICG increased after 30 minutes compared to 15 minutes in two sarcoma cell lines, and the MCF-7 breast cancer line (Figure 1a, 1d). At 40x magnification with differential interference contrast (DIC), the localisation of ICG uptake was shown to be cytoplasmic after 30 minutes incubation with 25μM ICG (Figure 1b). As the concentration of ICG increases the amount of intracellular uptake also increases, however at 50μM ICG, notable background signal was also detectable when HT1080 cells were incubated for 30 minutes (Figure 1c). Flow cytometry data using the HT1080 cell line showed that there is significant intracellular uptake at both 15 and 30 minutes and supports the findings on microscopy (Figure 1e).

**Figure 1.**
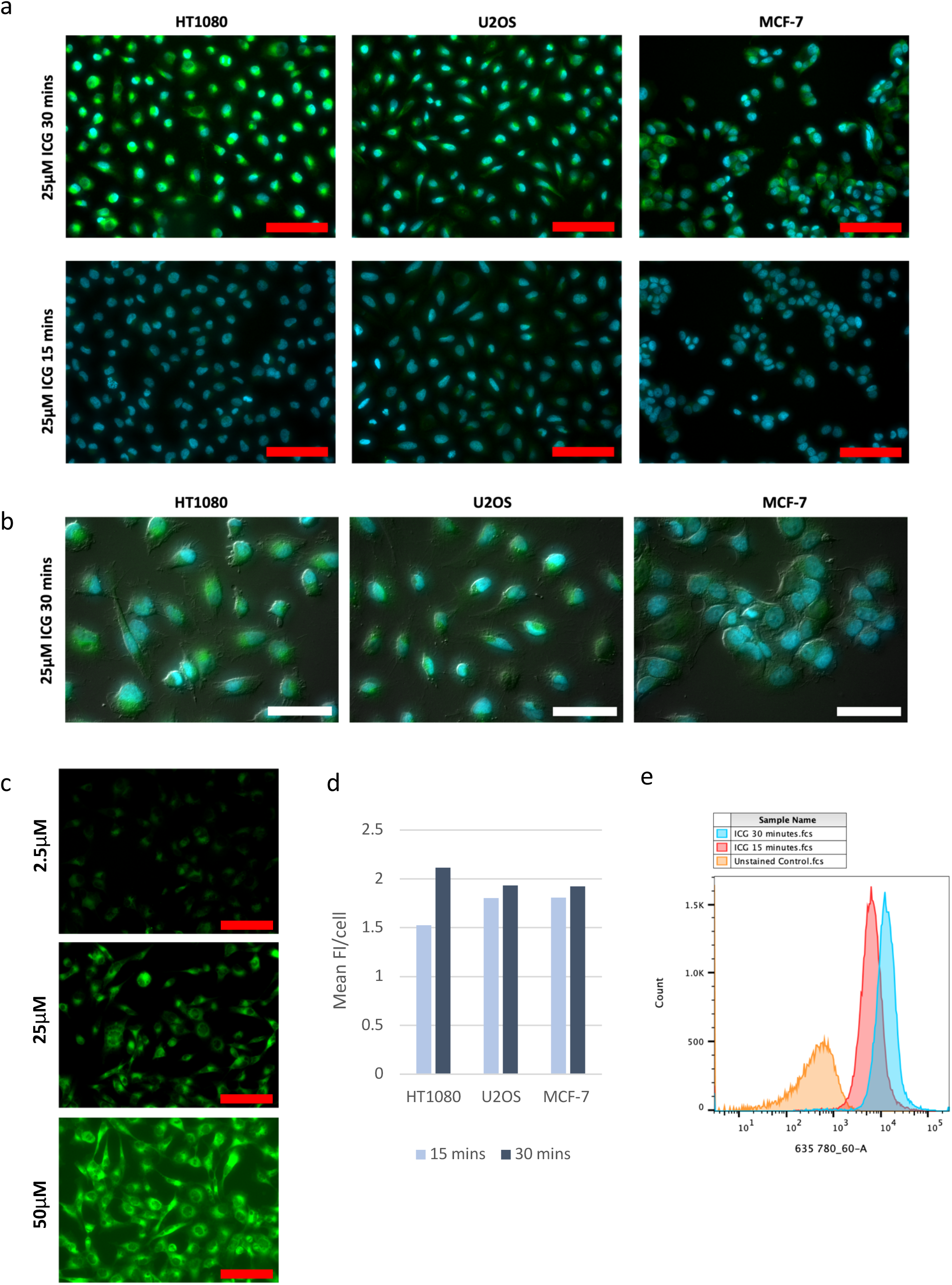
*In vitro* uptake of ICG in cancer cell lines at differing ICG concentrations and incubation times. (a) Fluorescent images of HT1080, U2OS and MCF-7 cells after incubation with 25μM ICG (green) counterstained with DAPI nuclear stain (blue) for 15 minutes vs 30 minutes (20x magnification). (b) Images of cell lines at 40x magnification after 30 minutes incubation with 25μM ICG (green), DAPI (blue) and DIC channel overlay. (c) HT1080 cells incubated with differing ICG concentrations for 30 minutes (2.5μM – 50μM); Cy7 channel only. (d) Graph comparing 15 mins vs 30 mins incubation of HT1080 cells with 25μM ICG using quantified data from 40x microscopy images. (e) Flow cytometry histogram data using HT1080 cells incubated with 25μM ICG for 15 minutes (red) and 30 minutes (blue) vs an unstained control (orange); ICG detected using laser 635 780/60-A. Red scale bars= 100μm and white scale bars= 50μm. Mean FI= mean fluorescence intensity, ICG= indocyanine green.

### Cellular ICG uptake correlates with cell line proliferation rate

The fluorescence intensity after 30 minutes incubation with 25μM ICG differed between the panel of cancer cell lines on both fluorescence microscopy imaging and flow cytometry. However, we observed a greater sensitivity for ICG detection using flow cytometry and therefore the comparison of inter-cell line differences in uptake was quantified using this method. A strong positive correlation was observed when plotting the median fluorescence intensity (MFI) against the proliferation rate (CCK-8) across the panel of cancer cell lines (Figure 2); Spearman’s rho correlation coefficient = 1.000, p<0.001. HT1080 cells, which are originally derived from a patient with an aggressive dedifferentiated chondrosarcoma and proliferate rapidly, exhibited the highest MFI when compared to other cancer cell lines. The least proliferative sarcoma cell line SAOS2 exhibited the lowest MFI after 30 minutes ICG incubation, alongside the MCF-7 breast cancer cell line which is widely considered to have low aggressiveness and low metastatic potential^44^.

**Figure 2.**
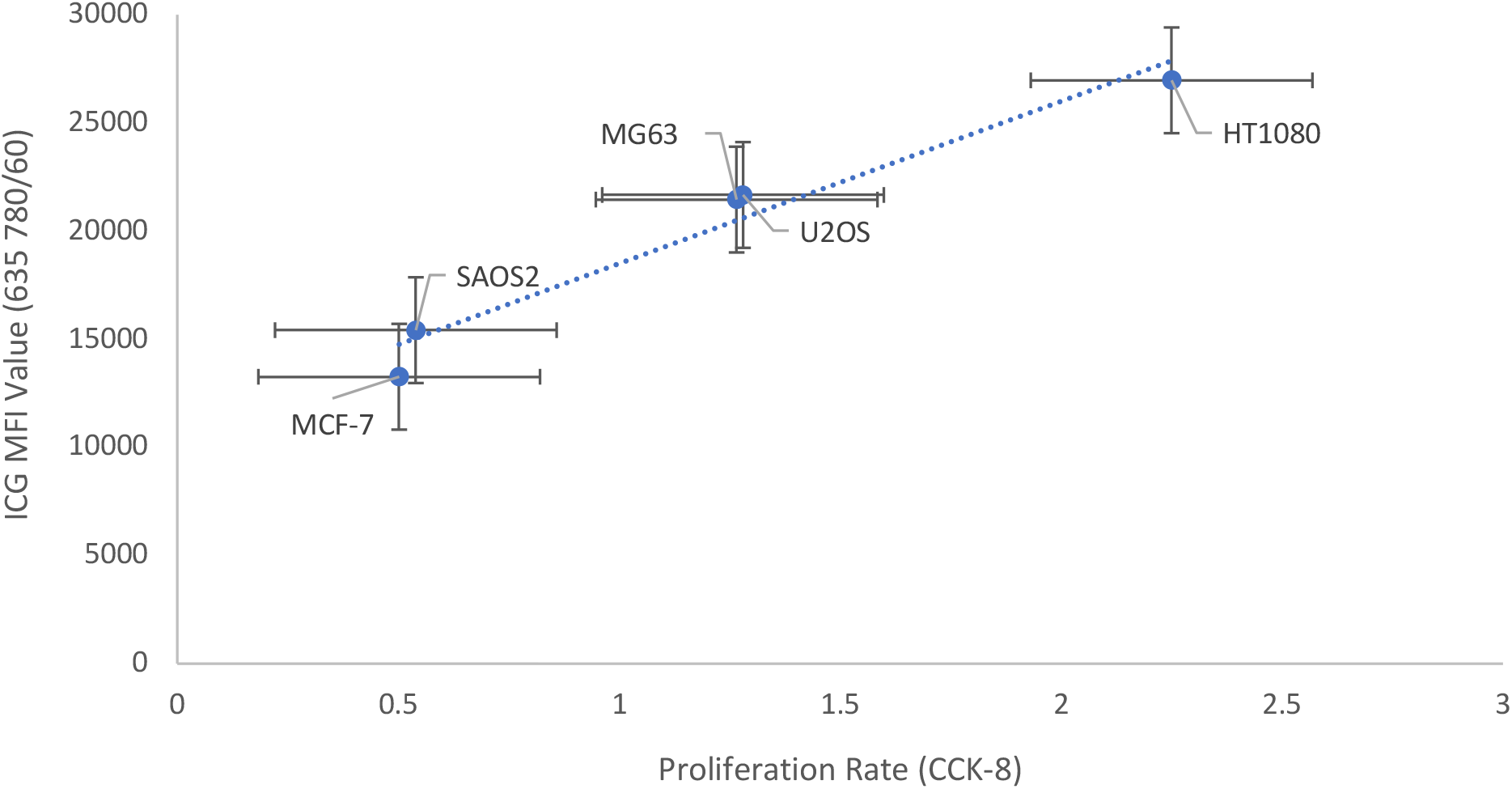
Scatter plot showing the correlation between flow cytometry median fluorescence intensity of ICG; 635 780/60 (y-axis) and cell line proliferation rate; CCK-8 (x-axis). Labelled data points indicate each cancer cell line, Mean ± SEM, with linear trend line (dotted blue). Correlation was calculated using SPSS software (V.27) non-parametric Spearman’s rho function; Rs = 1.000, p<0.001 (2-tailed). The analysed MFI and CCK data were taken as the mean value over three experiments, repeated on separate occasions. *MFI= median fluorescence intensity, CCK-8= cell counting kit-8, SEM= standard error of the mean*, ICG= indocyanine green.

### Retention of ICG after 24 hours varies between sarcoma cell lines

Given the differences in ICG cellular uptake between sarcoma cell lines, we investigated whether proliferation rate also correlated to the retention of cellular ICG. After 30 minutes incubation with 25μM ICG, followed by 24hrs in normal media, the residual MFI on flow cytometry followed a similar correlation (Figure 3a); Spearman’s rho = 0.9, p=0.037 for both 0.5hrs and 0.5hrs + 24hrs. However, the calculated retention percentage after 24hrs differed between cell lines (Figure 3b), and no longer correlated to the proliferation rate; Spearman’s rho = 0.3, p=0.624. The most proliferative sarcoma cell line (HT1080) had the highest retention rate after 24hrs (23.5%), whilst the lowest retention rate was observed in the U2OS cell line (8.39%) even though it is the second most proliferative cell line (Figure 3b). SAOS2 and MCF-7 cell lines both demonstrated low retention rates of ICG.

**Figure 3.**
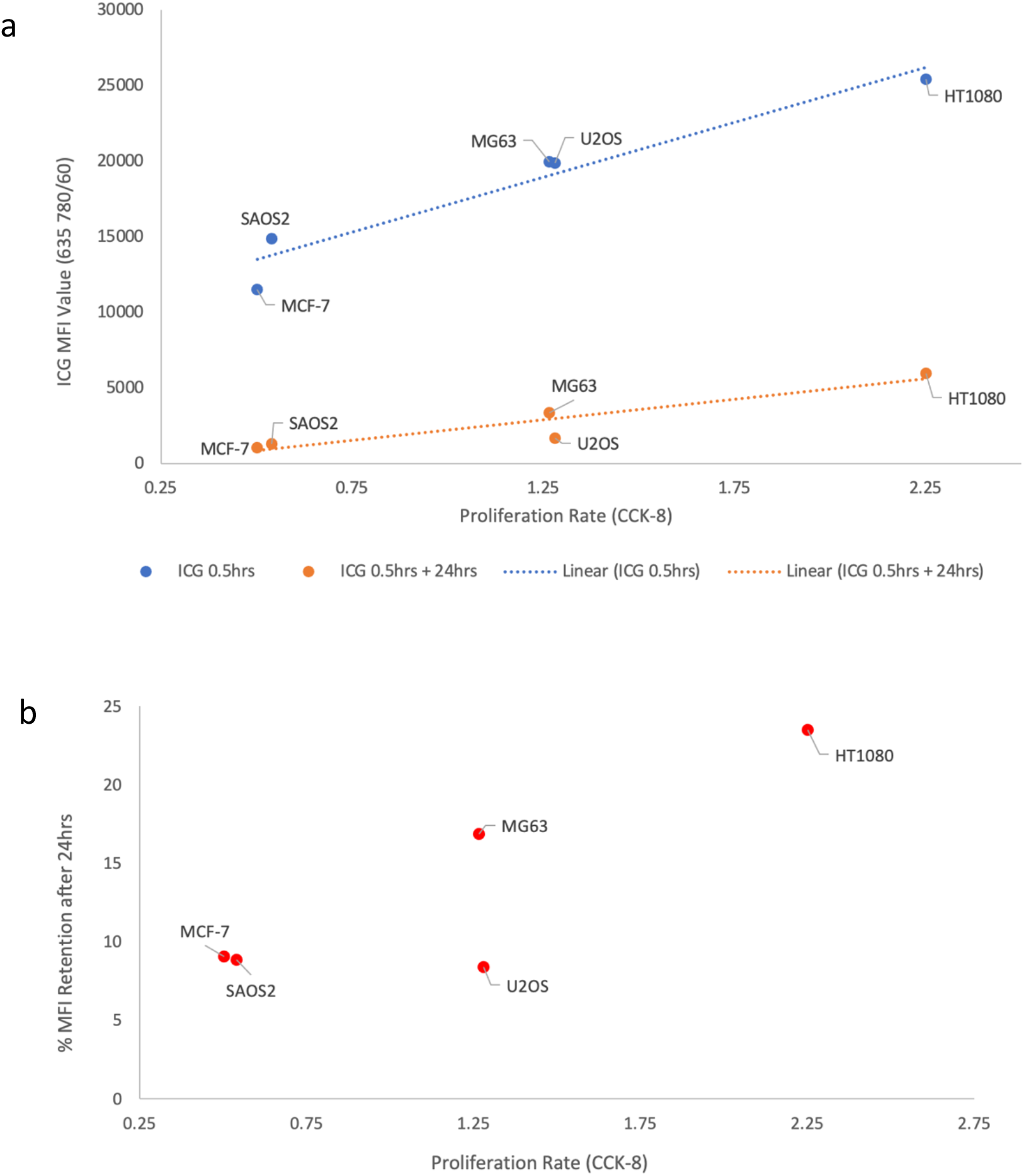
Scatter plots showing the correlation between ICG retention after 24hrs and cell line proliferation rate. (a) Scatter graph showing the correlation between MFI after 0.5hrs (blue) and 0.5hrs + 24hrs (orange) incubation in 25μM ICG vs proliferation rate (CCK) across all cell lines. The MFI data points were calculated as the average from two experiments repeated on separate occasions. Spearman’s rho function; Rs = 0.9, p= 0.037 for both data sets. (b) Scatter graph comparing ICG retention percentage to proliferation rate across all cell lines; Rs = 0.3, p=0.624. Flow cytometry data available in the Supporting Information. *MFI= median fluorescence intensity, CCK-8= cell counting kit-8*, ICG= indocyanine green.

### Pitstop 2 inhibits ICG uptake in sarcoma cells

When pre-incubated with PS2, a selective inhibitor of CME, subsequent uptake of ICG after 30 minutes was reduced. The level of inhibition was quantified with both flow cytometry (Figure 4a) and fluorescence microscopy (Figure 5b) using the HT1080 cell line.

**Figure 4.**
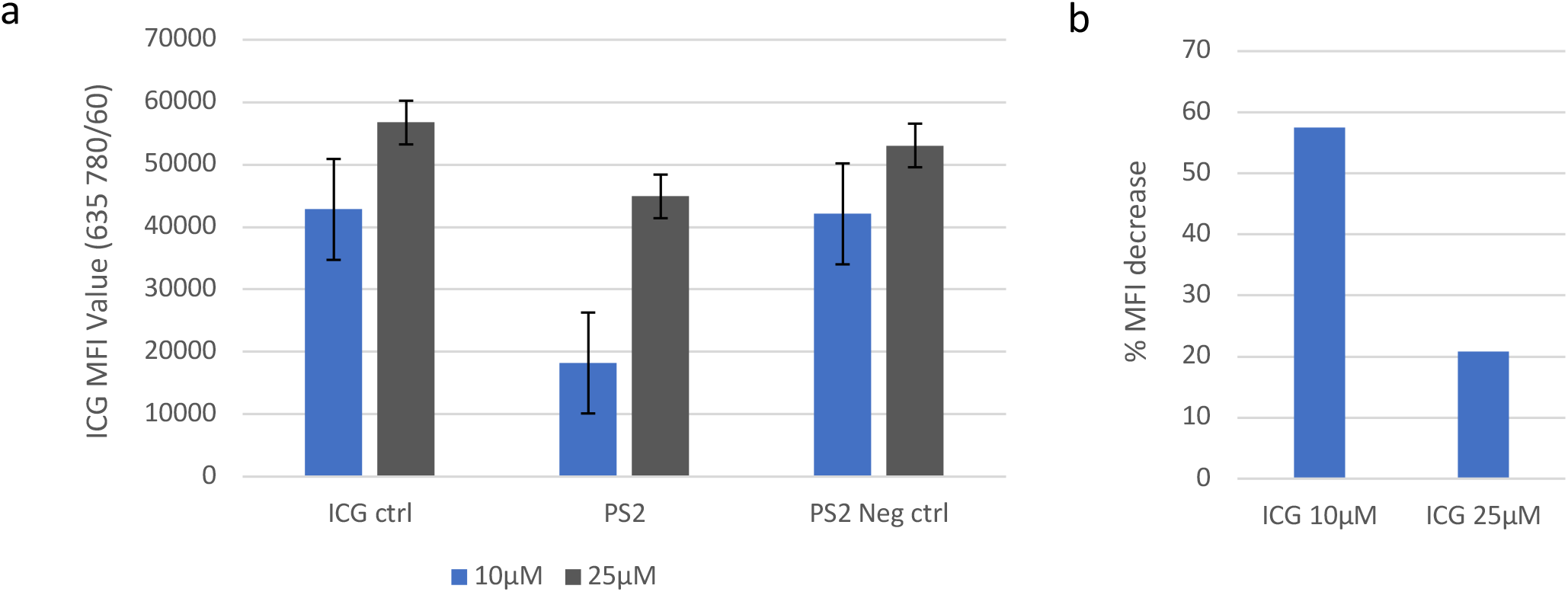
The effect of PS2 on the uptake of ICG in the HT1080 cell line at 10μM ICG vs 25μM ICG, incubated for 30 minutes. (a) MFI after treatment with 30μM PS2 inhibitor or 30μM PS2 negative control* followed by 30 minutes incubation with 10μM ICG (blue) or 25μM ICG (grey), compared to an untreated control. Mean ± SEM. (b) Percentage reduction in MFI observed in HT1080 cells after treatment with 30μM PS2 at differing ICG concentrations for 30 minutes. *MFI= median fluorescence intensity*, PS2= Pitstop 2. ICG= indocyanine green. *Pitstop 2 negative control compound has a highly related structure to Pitstop 2 but does not block receptor mediated endocytosis.

**Figure 5.**
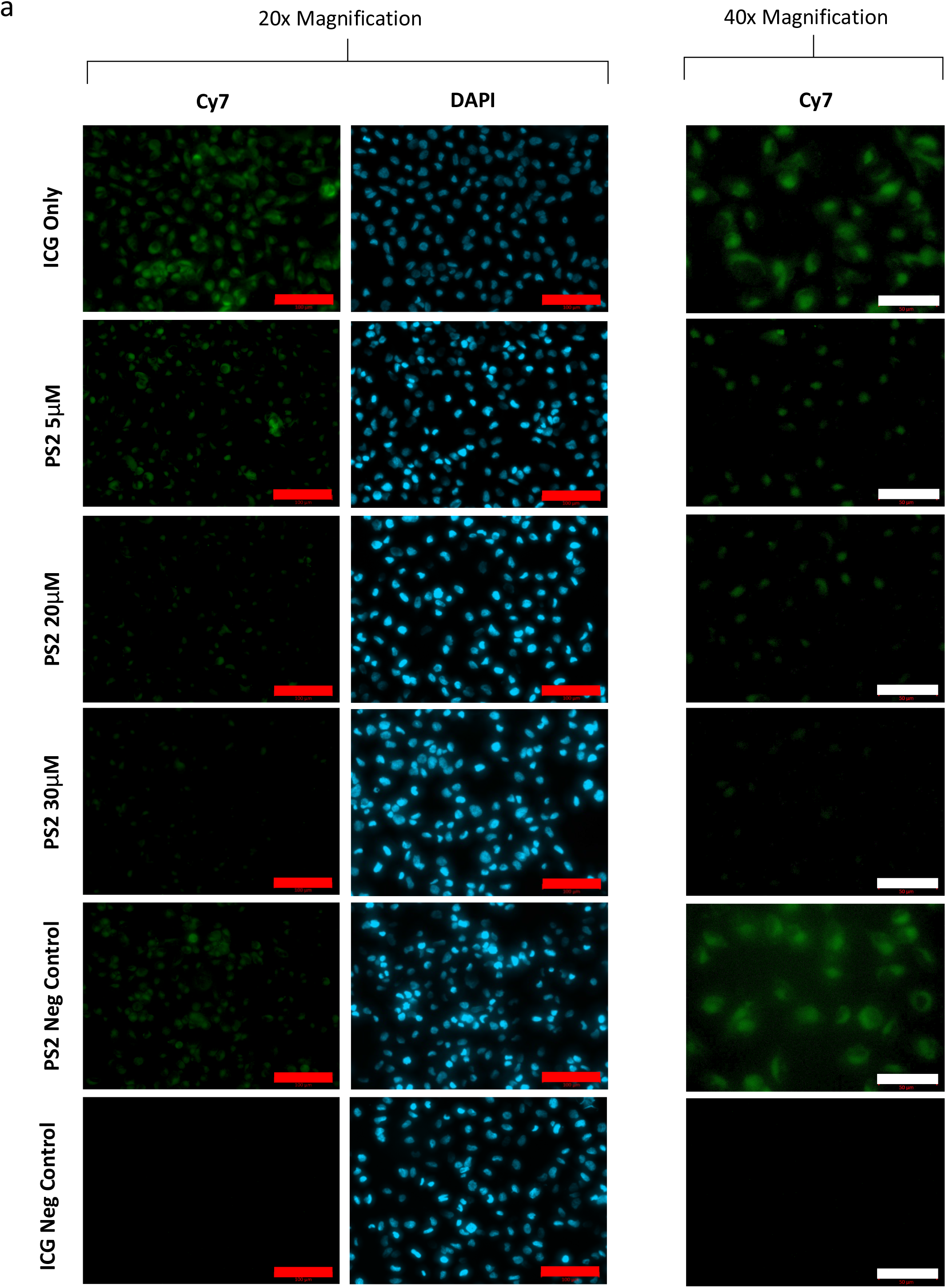
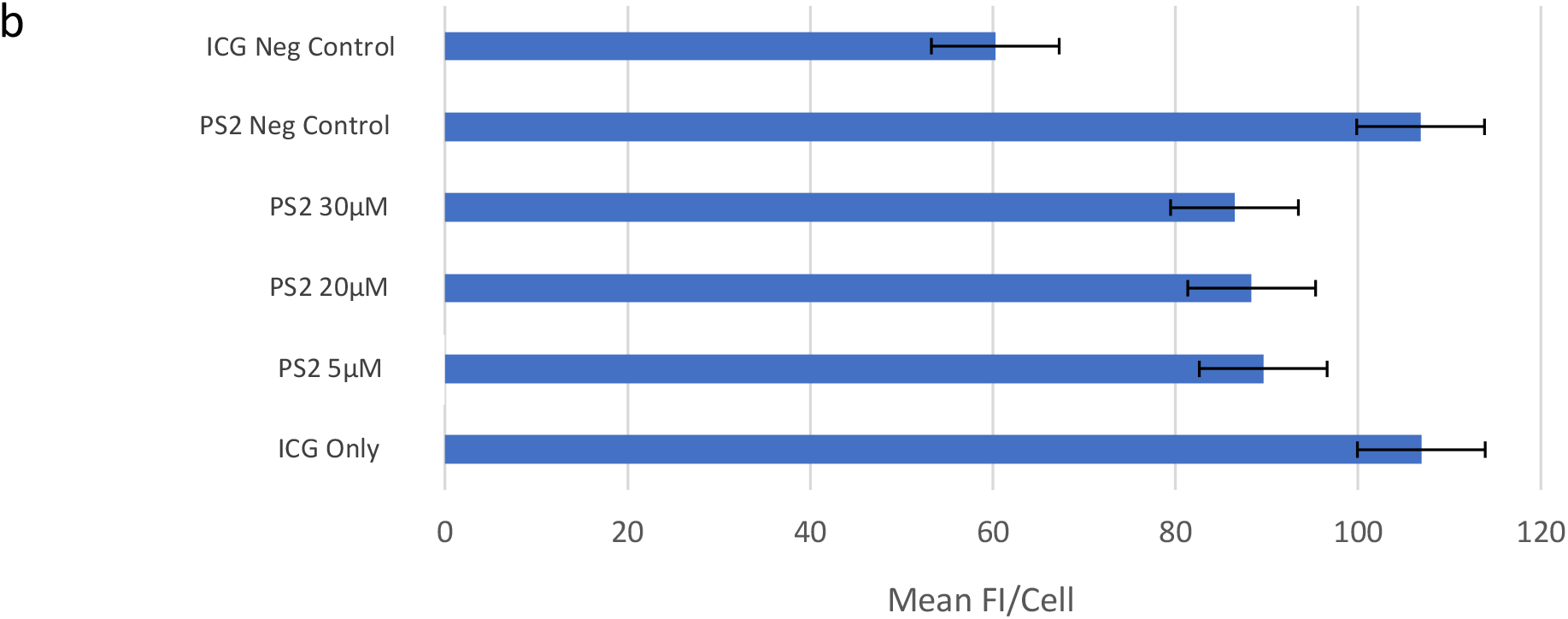
Data showing the effect of PS2 on ICG uptake in HT1080 cells at differing concentrations. (a) ICG uptake in HT1080 cells is inhibited by PS2 in a concentration dependent manner, when pre-incubated with 5μM, 20μM and 30μM of PS2 for 15 minutes followed by incubation with 25μM ICG for 30 minutes in the continue presence of the inhibitor. PS2 negative control* (30μM) does not inhibit ICG uptake. Images at 20x (left) and 40x (right) magnification show a decrease in intracellular ICG (green) at differing PS2 concentrations. Cy7 exposure time: 2000ms. DAPI exposure time: 80ms. Red scale bars = 100μm and white scale bars = 50μm (b). Quantification of fluorescence microscopy 40x images showing the effect of differing PS2 concentrations on the mean FI per cell. Mean ± SEM. Mean FI= mean fluorescence intensity, PS2= Pitstop 2, SEM= standard error of the mean, ICG= indocyanine green. *Pitstop 2 negative control compound has a highly related structure to Pitstop 2 but does not block receptor mediated endocytosis.

**Figure 6.**
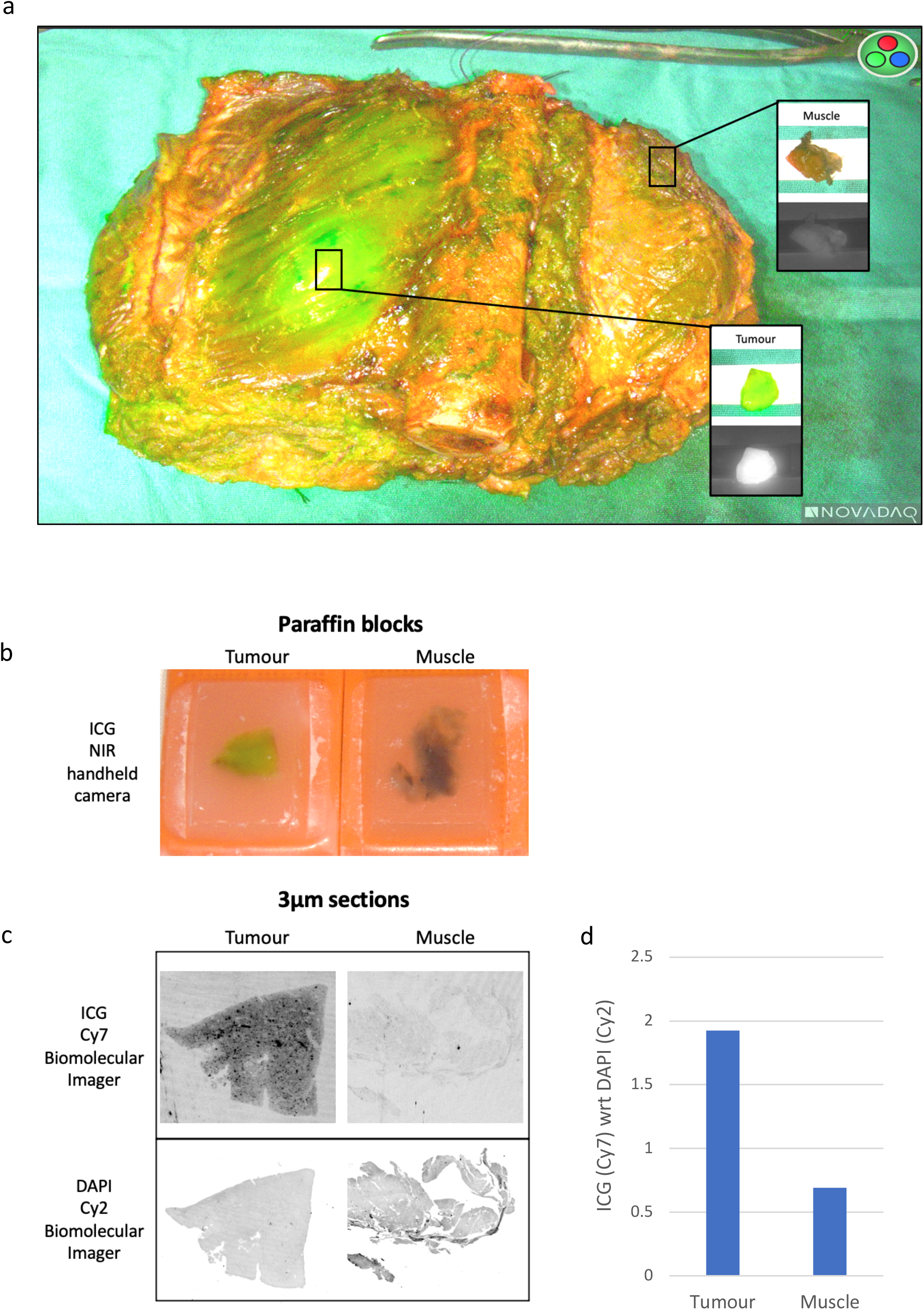
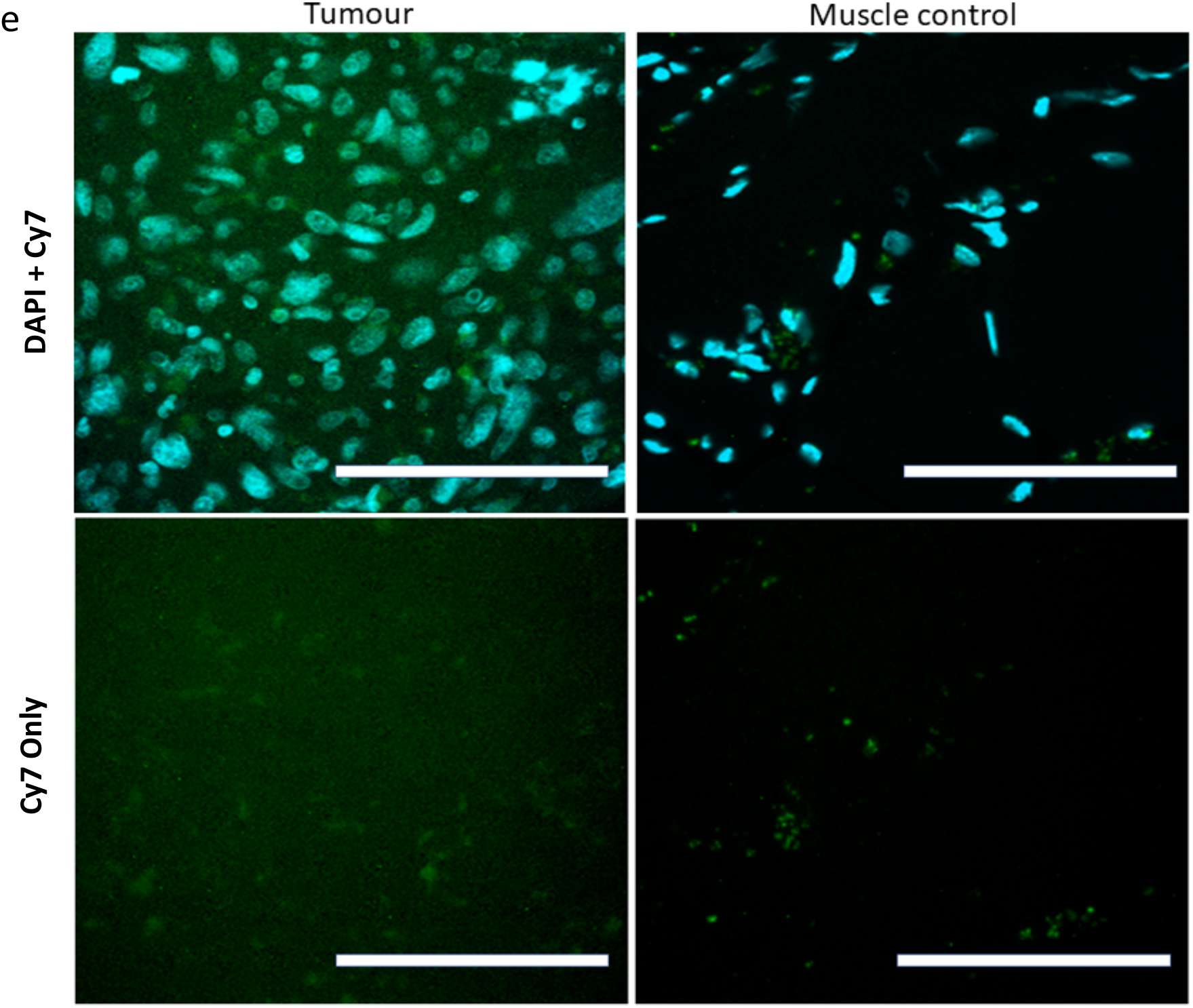
Images of ICG detection in human tumour samples vs non-tumour muscle control after administration of intravenous ICG pre-operatively. (a) Macroscopic images taken intraoperatively using the Stryker SPY-PHI handheld NIR camera in multiple camera modes. The resection was of a large grade 3 leiomyosarcoma specimen which had arisen in the anterior compartment of the thigh and required removal of a section of femur. Following delivery of the specimen, a tumour sample was taken from an area of intense fluorescence. A sample of muscle was taken to act as control tissue. (b) Specimens from ‘a’ were fixed in 4% formaldehyde and processed into paraffin blocks. Using the Stryker SPY-PHI handheld NIR camera, images were taken 18 days later of the blocks and demonstrated that fluorescence was still present indicating the ICG in the tumour was stable. (c) 3μm sections were imaged using the Amersham Typhoon Biomolecular Imager, again showing the presence of ICG in the tumour but not in the control, and the presence of DAPI in both specimens. (d) Tumour and muscle ICG (Cy7) signal quantified on Fiji Image J wrt DAPI (Cy2). (e) The same specimen slides from ‘c’ were imaged on the Zeiss Axio Imager 1 microscope, Cy7 exposure 15000ms. There was limited detection of ICG in the tumour cells, and some scattered ICG punctate signal in the muscle which appears to be extracellular. White scale bars= 100μm. NIR= near infrared. wrt = with respect to, ICG= indocyanine green.

From flow cytometry data, the relative level of PS2 inhibition appeared to be dependent upon the concentration of ICG (Figure 4a). Following treatment with PS2, when HT1080 cells were incubated for 30 minutes at lower concentrations of ICG (10μM), a greater reduction in MFI was observed when compared to higher ICG concentrations (25μM) (Figure 4b). The observed reduction in intracellular ICG uptake in the presence of PS2 was also greater at higher concentrations of the inhibitor (30μM) compared to lower concentrations (5μM) (Figure 5a, 5b). Although PS2 reduced the uptake of ICG compared to the PS2 negative control, it did not completely block ICG uptake to the level of the unstained control.

### ICG is detectable in human sarcoma tumour tissue vs muscle control

Following resection of the grade 3 leiomyosarcoma, ICG was visible in the human tissue specimen using the SPY-PHI NIR camera intraoperatively (Figure 6a). A sample of muscle tissue was sampled and did not fluoresce, acting as a negative control. The tissue samples were processed into paraffin blocks and images were re-taken 18 days later using the SPY-PHI NIR camera, demonstrating stability of ICG within the tumour (Figure 6b). After being processed into 3μm slides, imaging results from the Biomolecular Imager demonstrated significant ICG signal in the tumour slide compared to the muscle control (Figure 6c). On subsequent imaging of these slides at high magnification on the Zeiss Axio Imager 1, the detection of intracellular ICG was very limited, even at long Cy7 exposure times (15,000ms) (Figure 6e). There also appeared to be some non-specific aggregates of ICG within the muscle control. The quality of these microscopy images is poor and not suitable for quantification or comparative analysis, due to the limitation of our microscope’s efficiency within the Cy7 detection range.

## Discussion

The accumulation of ICG within tumours after intravenous administration has been reported across a number of cancer types^35, 37, 40, 45^. It is well known that the preferential tumour cell uptake of ICG is non-specific^45^. Although several potential mechanisms have previously been suggested and explored, including tight junction disruption, the expression of specific membrane transporters, and dysregulated endocytic pathways, the exact mechanisms are still not fully understood^33, 40^. It is appreciated that the *in vivo* pharmacokinetics of ICG uptake in tumours will differ greatly to *in-vitro* studies, as factors such as serum protein binding, tumour vascularity, and the enhanced permeability and retention (EPR) effect will influence the uptake and availability of ICG to tumour cells. The timing of administration of ICG preoperatively will also have a large effect on tumour retention and will effect the signal to background ratio at the time of resection. The EPR effect in particular has been suggested to play an important role in tumour accumulation of ICG compared to healthy tissue, however other studies, as previously mentioned, have suggested that passive tumour cell targeting of ICG and preferential ICG uptake and retention by tumour cells also has a significant role^40^.

Our data contributes to previous studies and suggests that the rate of ICG uptake differs depending on the cancer cell type, with more proliferative cell lines having increased uptake within 30 minutes^40^. However, the cellular retention of ICG after 24hrs differed between sarcoma cell lines and, unlike the rate of ICG uptake, did not correlate to proliferation rate. This supports the aforementioned concept of passive tumour cell targeting by ICG via increased uptake, however it suggests that the retention of ICG within a tumour is likely to be highly dependent upon the EPR effect. In the U2OS cell line for example, we observed a high uptake rate of ICG, whilst the retention rate after 24hrs was the lowest across our panel of cell lines.

We observed a significant correlation between ICG cellular uptake and cell line proliferation rate. The proliferation capacity of a cancer cell line is often used as an indicator of “aggressiveness”, and such cell lines often exhibit marked genetic alterations, dysregulated cellular pathways and the presence of activated oncogenes which contribute to high invasive potential^44^. From our study, given the relatively short 30 minutes ICG incubation time, with comparable cell numbers at the time of incubation, the results observed are unlikely to be due to differences in the number of cells. The preferential uptake of ICG by the more “aggressive” cell lines can therefore be postulated to be due to differences in intrinsic cell mechanisms, such as dysregulated clathrin mediated endocytosis (CME) and endosome trafficking.

Our data suggest that the primary mechanism of ICG uptake in sarcoma cells is by CME^40^. In this study, PS2, a novel cell-permeable clathrin inhibitor, reduced the uptake of ICG in a concentration dependent manner, suggesting that this is the principal method of ICG uptake in sarcoma cells. However, our data showed that at increasing ICG concentrations, the effect of PS2 inhibition was reduced. This suggests that other endocytic mechanisms such as caveolae-dependent endocytosis may also play a role, but at lower concentrations of ICG cells are largely dependent on CME for effective ICG uptake. It is also important to mention that some recent studies have demonstrated PS2 to also inhibit clathrin independent endocytosis, suggesting only partial selectivity of this compound for CME^46, 47^.

Following the surgical resection of a grade 3 leiomyosarcoma, ICG was detectable within the tumour macroscopically and also present on 3μm tumour sections. The ICG within this specimen was also detectable in the paraffin embedded tumour blocks 18 days later. This suggests high stability of ICG within fixed tumours for extended periods of time. One of the limitations to this study was the poor detection of ICG at the cellular level in the human sarcoma tissue sections, even at long exposure times. We hypothesise that the concentration of ICG per cell in these slides is low, and as such postulate that ICG is in its monomeric form^5^. This likely results in a far red detection range (excitation >780nm) which exceeds the capability of our current set-up, which has an emission quantum efficiency of <5% in the Cy7 channel. For the *in vitro* cell line studies, the concentration of ICG available per cell will be far greater than *in vivo*, allowing suitable detection with our current equipment, due to the spectral properties of ICG in oligomer form^5^. Currently, this work is limited by the lack of availability of microscopes with efficient far red and NIR capabilities.

## Conclusion

This study found that cellular ICG uptake correlated significantly with cell line proliferation rate across a panel of sarcoma cell lines, whilst ICG retention did not. From this study, it can be suggested that the use of ICG for tumour detection in sarcoma surgery may demonstrate higher utility in high grade tumours compared to low grade tumours, due to enhanced uptake at the cellular level in the more aggressive cell lines. It is also likely that the EPR effect plays a significant role in the retention of ICG within tumours. Although further work is required to understand the exact mechanisms causing enhanced ICG uptake in more proliferative cell lines, our data supports CME as the principle endocytic pathway for ICG uptake. Future work on the detection of ICG at the cellular level within human tissue sections is required, with the aid of purpose-built NIR microscopes. This will be an important step to help guide the use of ICG in clinical practice for sarcoma surgery and to maximise its translational benefit in the future.

## Acknowledgements

The authors would like to thank Newcastle University BioImaging Facility for their help and support with this research project.

