## Supporting Information for "Investigating the mechanisms of indocyanine green (ICG) cellular uptake in sarcoma"

**Cell Culture**

All cell lines were cultured in RPMI 1640 medium supplemented with 10% foetal bovine serum, 100 U penicillin/mL and 0.1 mg streptomycin/mL; incubated at 37°C in a 5% $CO_2$ humidified atmosphere.

For experiments involving PitStop2, the media used was DMEM/F12 (1:1) with 15mM HEPES, supplemented with 100 U penicillin/mL and 0.1 mg streptomycin/mL with no added foetal bovine serum.

| CELL LINE | ORIGIN | DOUBLING TIME (HOURS) |
|---|---|---|
| **HT1080** | ATCC | 26 |
| **U2OS** | ATCC | 29 |
| **SAOS2** | ATCC | 43 |
| **MG63** | A gift from Dr Neil Cross, Sheffield University, UK | 28 |
| **MCF-7** | ATCC | 30-40 |

Table S1. Information on the cell lines used in this study. ATCC= American Type Culture Collection.

**Flow Cytometry**

*Flow Buffer Formulation:* 500 mL PBS, 2.5 mL of 0.2 mmol/L EDTA prepared from anhydrous stock (Sigma), 2.5 mL of BSA Stock Solution (MACS 10% BSA).

Flow cytometric analysis of cellular ICG fluorescence was performed on the BD FACSCanto II Cell Analyser, using the 635 nm laser with bandwidth 780/60. Samples were analysed at a low flow rate for 100,000 total events. Flow cytometry data was analysed using FlowJo V10 Software. The main cell population was gated, doublets discriminated using FSC-H and a histogram with log scale generated against 635;780/60-A. The median fluorescence intensity (MFI) for each cell line was calculated by (MFI ICG Positive Sample – MFI Unstained Control Sample). MFI values were obtained using the statistical function within FlowJo V.10 software.

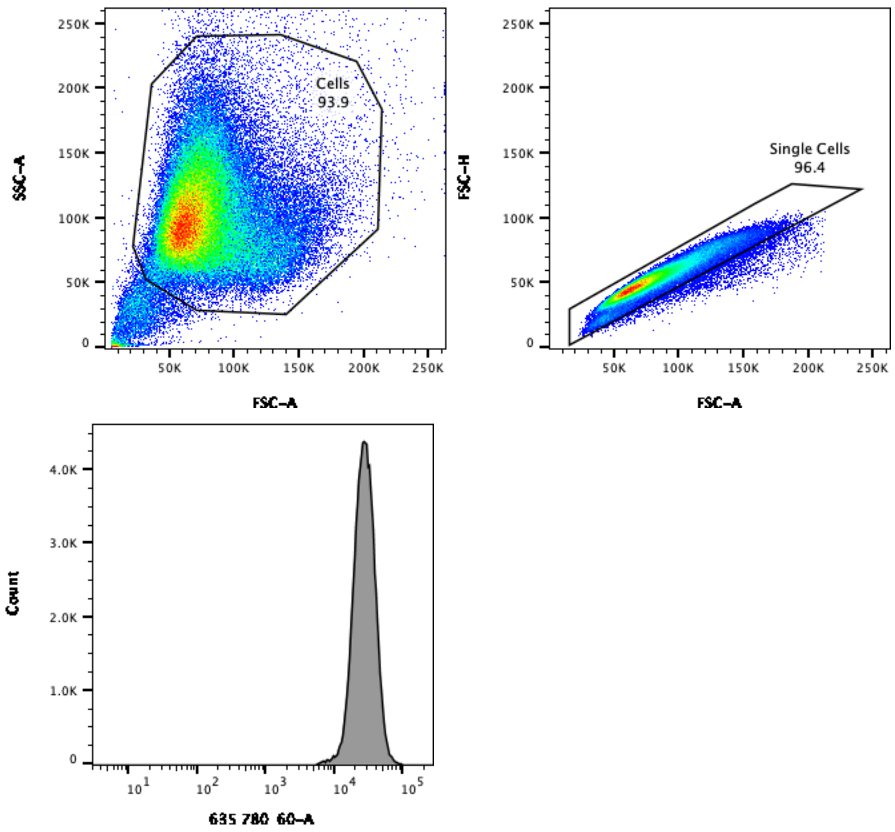


**Figure S1.** Demonstration of how how FACS data was processed for each sample. Main cell population was gated on FSC-A/SSC-A, doublets were disriminated via gating on FSC-A/FSC-H, and histogram of intensity of 635 (780/60) generated (x-axis log scale). Analysis performed on FlowJo V.10 software.

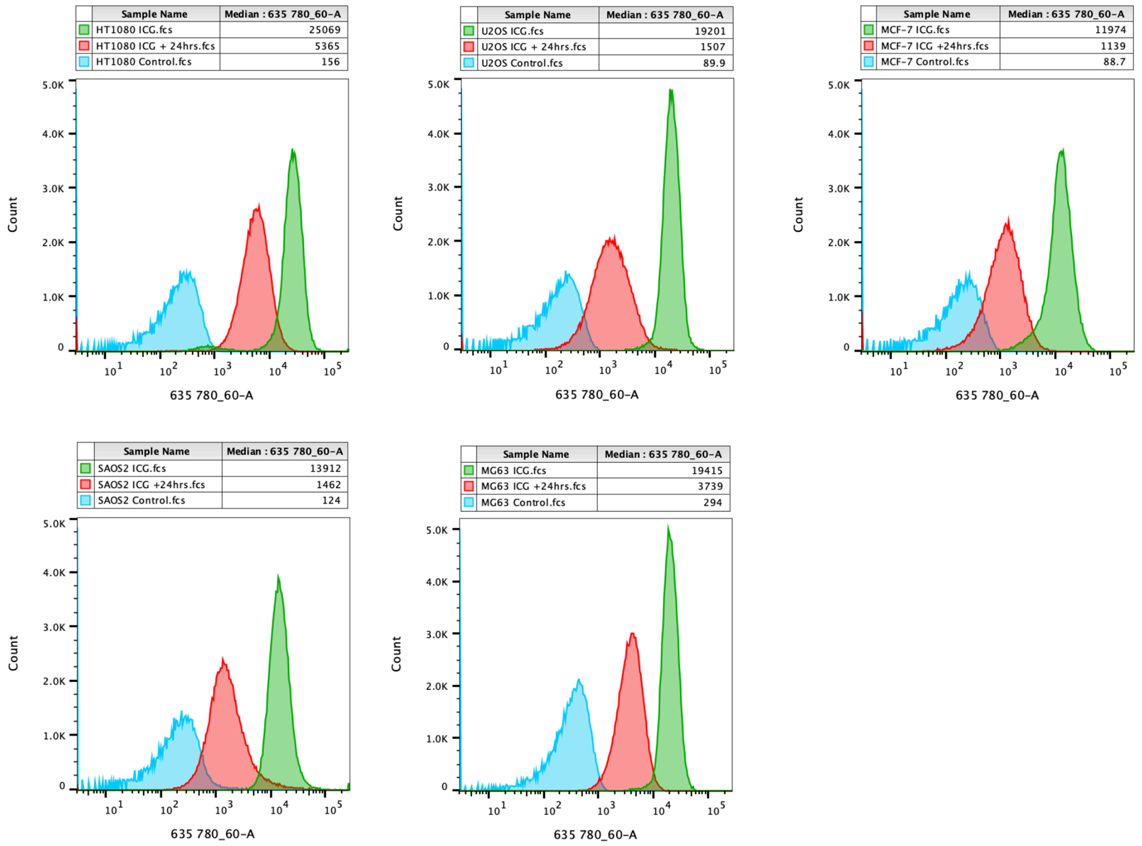


**Figure S2.** Example of FACS data generated across all cell lines for ICG uptake within 30 minutes, followed by normal media for 24hrs to determine ICG uptake and retention rate. 25µM ICG concentration. Data analysed using FlowJo V10 Software.

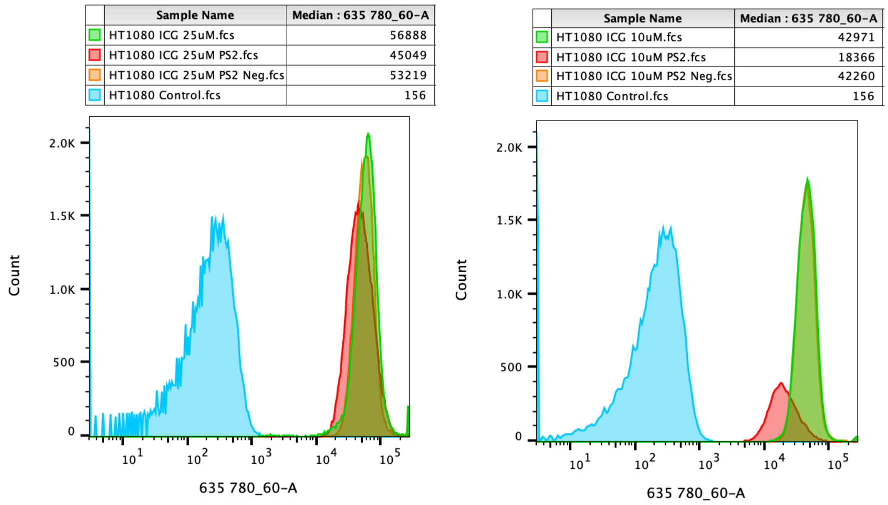


**Figure S3.** Flow cytometry data showing the effects of PS2 treatment on ICG uptake at 25µM and 10µM ICG. Experiment performed on the HT1080 cell line, pre-treated with 30µM PS2 or 30µM PS2 negative control for 15 minutes, followed by incubation with ICG for 30 minutes in the continued presence of the inhibitor. Data analysed using FlowJo V10 Software.

**Fluorescence Microscopy**

Microscope: Zeiss Axio Imager 1.
Camera: Zeiss MRm (Monochrome). Light Source: HXP120 metal halide, 825uW +/- 1.65uW.
Semrock Cy7-B cube bandpass filters: ex 670-745, em 768-850 (Exposure 2000ms – 15,000ms, light source intensity 100%). DAPI/Zeiss 49 Cube bandpass filters: ex 335-385, em 420-470 (Exposure 80ms, light source intensity 100%).
Zeiss Lenses: 20x Plan-Apochromat 20x/0.8 M27; 40x EC Plan-Neofluar 40x/0.75 M27.
All fluorescent images taken with NIR mode active and 1x1 binning.
DIC light source: TL VIS-LED Lamp (light source intensity 10%), exposure time 400ms.

**Image Quantification using Zen 3.3 Blue Edition Software (Zeiss)**

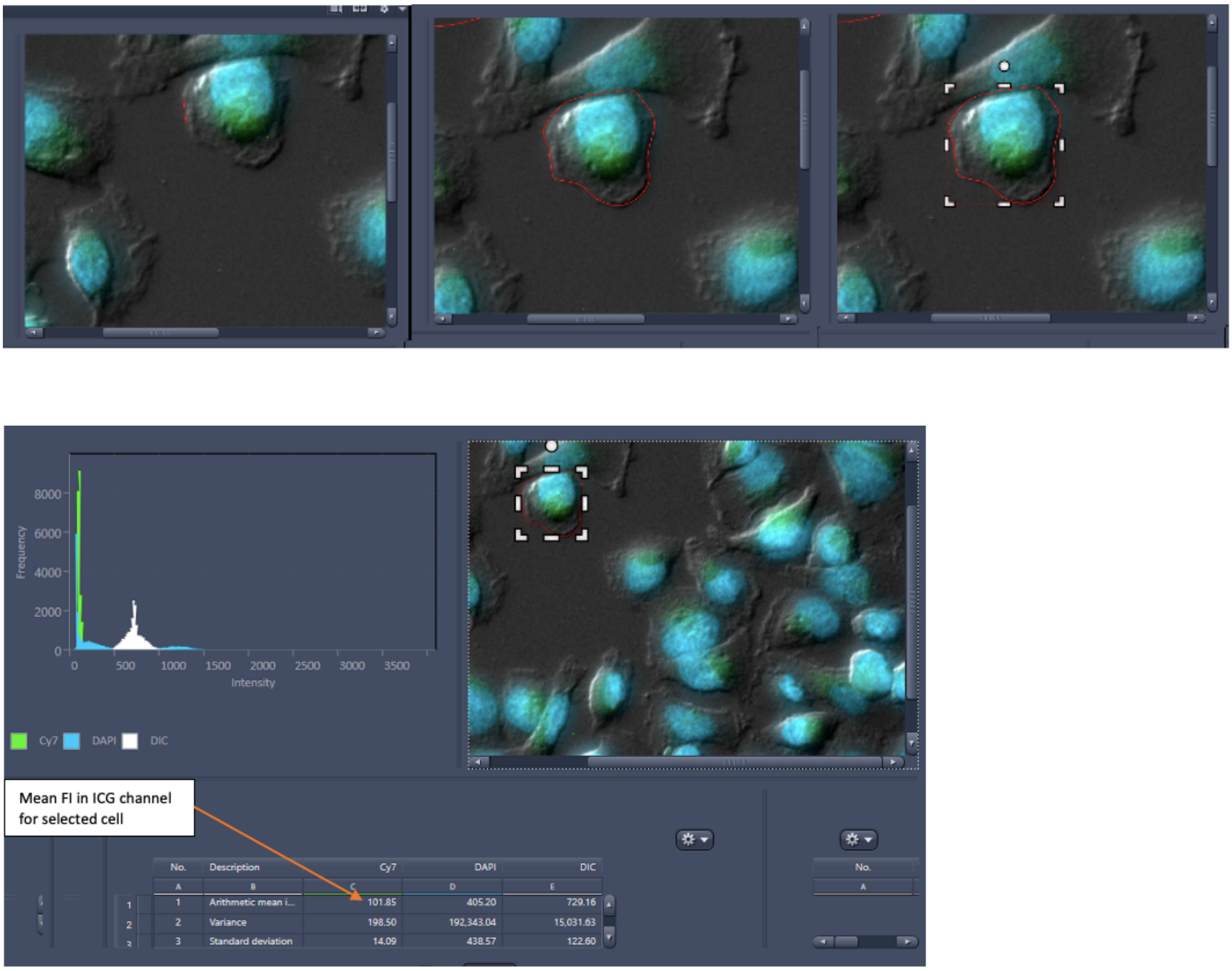


**Figure S4.** Zen software polygonal tool was used to draw around each cell at 40x magnification using DIC to guide the identification of individual cells. The mean florescence intensity (MFI) per cell was generated within the Cy7 column of the table on the histogram image view tab. Four representative cells were analysed per FOV with this method, and the average MFI calculated. Screenshots taken from ZEN 3.3 Blue Edition software (Zeiss).
